# RanBALL: An Ensemble Random Projection Model for Identifying Subtypes of B-Cell Acute Lymphoblastic Leukemia

**DOI:** 10.1101/2024.09.24.614777

**Authors:** Lusheng Li, Hanyu Xiao, Xinchao Wu, Zhenya Tang, Joseph D. Khoury, Jieqiong Wang, Shibiao Wan

## Abstract

As the most common pediatric malignancy, B-cell acute lymphoblastic leukemia (B-ALL) has multiple distinct subtypes characterized by recurrent and sporadic somatic and germline genetic alterations. Identifying B-ALL subtypes can facilitate risk stratification and enable tailored therapeutic design. Existing methods for B-ALL subtyping primarily depend on immunophenotyping, cytogenetic tests and genomic profiling, which would be costly, complicated, and laborious. To overcome these challenges, we present **RanBALL** (an ensemble **Ran**dom projection-based model for identifying **B**-**ALL** subtypes), an accurate and cost-effective model for B-ALL subtype identification. By leveraging random projection (RP) and ensemble learning, RanBALL can preserve patient-to-patient distances after dimension reduction and yield robustly accurate classification performance for B-ALL subtyping. Benchmarking results based on >1700 B-ALL patients demonstrated that RanBALL achieved remarkable performance (accuracy: 0.93, F1-score: 0.93, and Matthews correlation coefficient: 0.93), significantly outperforming state-of-the-art methods like ALLSorts in terms of all performance metrics. In addition, RanBALL performs better than tSNE in terms of visualizing B-ALL subtype information. We believe RanBALL will facilitate the discovery of B-ALL subtype-specific marker genes and therapeutic targets to have consequential positive impacts on downstream risk stratification and tailored treatment design. To extend its applicability and impacts, a Python-based RanBALL package is available at https://github.com/wan-mlab/RanBALL.

## Background

B-cell acute lymphoblastic leukemia (B-ALL) is a hematological malignancy that originates from the precursor B-cells of the bone marrow. As the most common type of acute lymphoblastic leukemia (ALL), B-ALL accounts for approximately 5,000 cases in the United States each year, especially affecting children under the age of five (1,2). The clinic diagnostic and biologic heterogeneity of B-ALL present a significant challenge in terms of subtype classification and risk stratification (3,4). Previous studies have highlighted the necessity of precise subtype identification for highly diverse therapeutic approaches for each patient (5–8). So far, multiple distinct B-ALL subtypes have been characterized through recurrent and sporadic somatic and germline genetic alterations, e.g., BCR-ABL1 (Philadelphia (Ph) chromosome), TCF3-PBX1 (9), hypodiploid (10), etc. Based on integrated genomic analysis of 1,988 childhood and adult cases, Gu et al. (11) identified 23 B-ALL subtypes by chromosomal rearrangements, sequence mutations and heterogeneous genomic alterations. In addition, another study (12) comprehensively reviewed the etiologic heterogeneity of childhood acute lymphoblastic leukemia (ALL) across different subtypes, highlighting the critical need for further investigations into risk factors that are specific to each subtype.

Conventional methods for B-ALL subtype identification primarily depend on a combination of morphological, immunophenotypic, cytogenetic, and molecular characteristics (13,14). With advancements in the next-generation sequencing (NGS) (15,16), RNA-seq has become an effective tool to unveil chromosomal rearrangements in individual tumors for genetic or clinical marker discovery (11,17). It has been increasingly investigated as an advanced diagnostic method for ALL clinical trials, underscoring its capability to accurately diagnose specific molecular subtypes (18,19). In addition, large cohort studies for new subtype detection and rapid classification with large-scale datasets raise more interest in the progress of the precision medicine (20–22). Categorizing new B-ALL subtypes typically involves integrating different types of NGS techniques such as whole-genome sequencing (WGS) (23), whole-exome sequencing (WES) (24), cytogenetic assays (25), etc. However, these methods for subtype identification would be costly, complicated, and time-consuming in clinical applications. In addition, extensive manual review of the results from these analyses like fusion detection and mutation calling is required, which is a laborious process.

Recognizing these challenges, the application of machine learning (ML) models to B-ALL subtype identification has emerged as a promising approach to revolutionize our understanding of this disease and improve patient outcomes (26). In recent years, multiple subtype classification tools based on different ML algorithms have been introduced based on well-defined B-ALL subtypes from WHO-HAEM5 (27), and ICC (28) classification system. For instance, Allspice (17) was developed to predict the B-ALL subtypes and driver genes based on a centroid model. However, centroid methods in Allspice may not fit in scenarios involving overlapping classes or imbalanced datasets. In addition, ALLSorts (29) was introduced to classify 18 B-ALL subtypes with logistic regression, which nevertheless could perform poorly as the relationships between features and subtypes were more complicated than a simple logistic model. Furthermore, ALLCatchR (30) was proposed based on integrating linear support vector machine (SVM) and gene set-based nearest-neighbor models. However, nearest-neighbor models in ALLCatchR relied heavily on calculating distances between data points, which could be computationally expensive for large datasets, especially as the number of samples or features increased. Recently, MD-ALL (31) was developed to predict B-ALL subtypes using SVM and PhenoGraph algorithms. Nevertheless, PhenoGraph’s clustering relied on the number of nearest neighbors to construct a graph, which might significantly affect classification results.

To address these concerns, we presented **RanBALL** (an Ensemble **Ran**dom Projection-Based Model for Identifying **B**-Cell **A**cute **L**ymphoblastic **L**eukemia Subtypes), an accurate and cost-effective model for B-ALL subtype identification based on transcriptomic profiling only. By leveraging the RP, RanBALL transformed the high-dimensional feature of gene expression data to lower-dimensional representations, effectively minimizing redundant and irrelevant information. To obtain robust performance of RanBALL model, multiple RPs were applied to the original gene expression matrix. The resulting dimension-reduced matrices were then used for an ensemble of SVM classifiers to generate the final subtype predictions. Benchmarking results based on a comprehensive B-ALL dataset demonstrated that RanBALL achieved significantly and consistently better performance than state-of-the-art B-ALL subtyping methods in term of accuracy, F1-score and Matthews correlation coefficient. We anticipate that RanBALL will bring substantially positive and direct impacts on clinical diagnosis improvement, risk stratification, and the development of personalized treatment strategies for B-ALL patients.

## Methods

### B-ALL datasets

The RNA-seq data and clinical information of B-ALL samples were obtained from St. Jude Cloud (https://pecan.stjude.cloud/static/hg19/pan-all/BALL-1988S-HTSeq.zip). The dataset included 1988 samples that were classified as 23 B-ALL subtypes from the study (11). In data processing, samples with two subtypes and those identified as “other” categories were filtered out. Additionally, the samples were processed by referring to the classification architecture outlined in the ALLSorts classifier (29). Due to the limited number of samples in subtypes “ZNF384-like” and “KMT2A-like”, "ZNF384-like" samples were merged with "ZNF384" samples to form a broader category labeled the "ZNF384 Group", and "KMT2A-like" samples were merged with "KMT2A" samples to form a broader category labeled the "KMT2A Group". Samples classified as the “CRLF2(non-Ph-like)” subtype were excluded due to limited sample size. After filtering, the B-ALL dataset contained a total of 1,743 samples across 20 distinct categories (Fig. 1A). The breakdown of the B-ALL dataset among the various subtypes was depicted in the pie chart (Fig. 1A). This B-ALL dataset was very imbalanced, with a few categories (such as Ph-like, High hyperdiploid and ETV6-RUNX1) taking up a significantly larger proportion (i.e., 18.6%, 15.4% and 10.8%) of the total compared to others. As illustrated in the Fig. 1B, the majority (about 81.0%) of patients for the B-ALL dataset belonged to the childhood and AYA groups. The distribution showed a higher concentration of B-ALL cases in younger age groups, with notable peaks in childhood and young adulthood.

**Figure 1.**
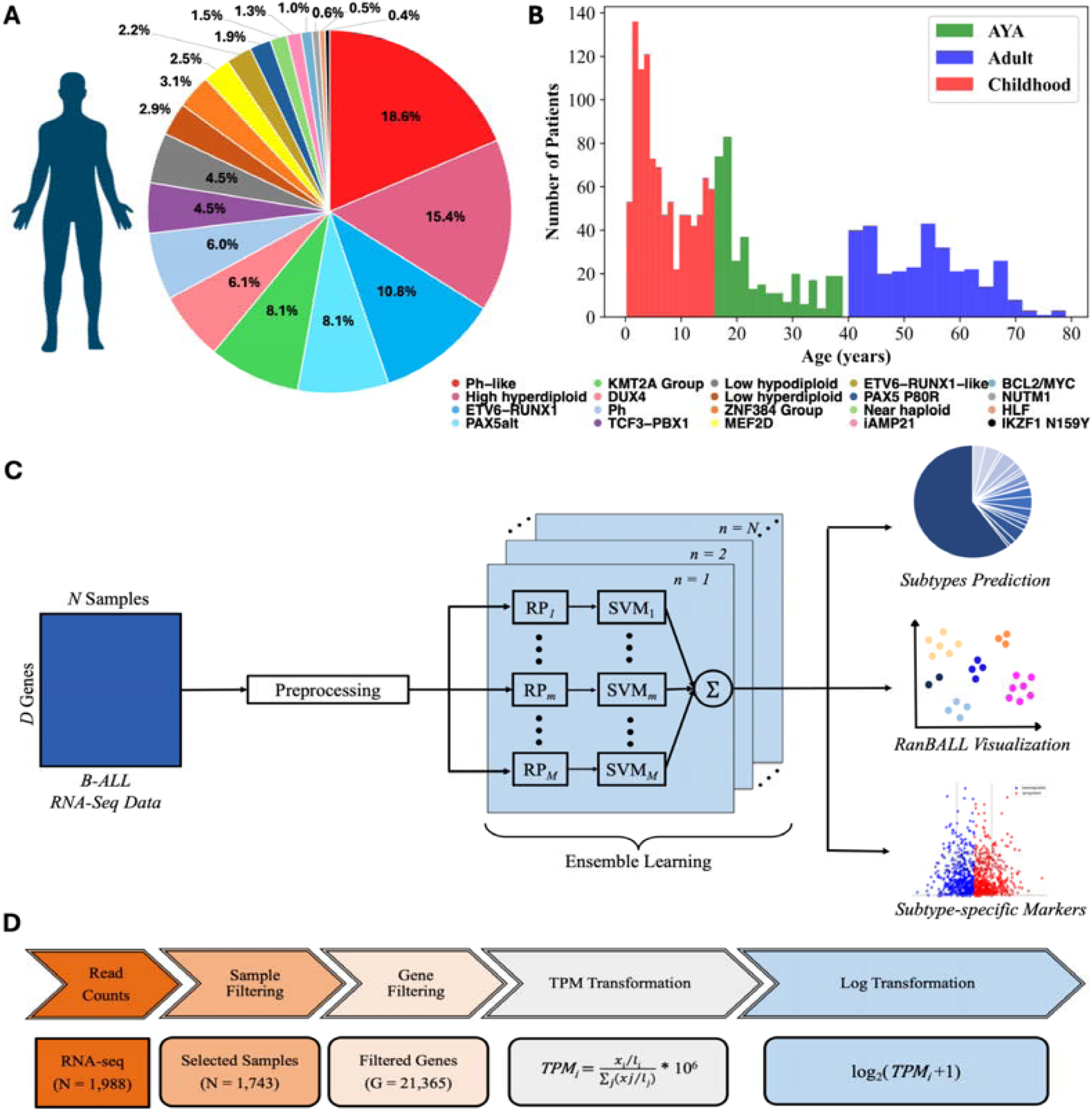
Overview of B-ALL subtype identification study using RanBALL framework. **(A)** The breakdown of the B-ALL dataset. The pie chart showed the distribution of 1,743 B-ALL samples across 20 molecular subtypes, each represented by a distinct color. Percentages reflected the relative prevalence of each subtype within the dataset. **(B)** The age distribution of the B-ALL dataset. The histogram illustrated the number of patients within each age group across three categories: childhood (red), adolescent and young adult (AYA, green), and adult (blue). **(C)** The framework of RanBALL. The feature dimension of preprocessed data was reduced by RP, and an ensemble of SVM classifier was trained on multiple dimensionally reduced matrices. In this framework, the dimensionality to be reduced to was predefined as 1200. The symbol *m* represents the *m*-th reduced-dimensional data matrix, while *n* denotes the predicted subtype. The RanBALL framework was designed to classify distinct subtypes, with the final prediction obtained through an aggregated output from the ensemble. Beyond subtype prediction, RanBALL also facilitated enhanced visualization of subtype clusters and the identification of subtype-specific markers, providing additional insights into the biological characteristics of each subtype. **(D)** Data preprocessing pipeline. The flowchart outlines the multi-step preprocessing applied to the RNA-seq data, starting with raw read counts and ending with log-transformed TPM values for 21,365 genes from 1,743 selected samples.

### RanBALL framework

RanBALL is an ensemble random projection-based multi-class classification model specifically designed for B-ALL subtyping using gene expression profiling. Leveraging the RP and SVM techniques, our model accurately and efficiently identifies distinct B-ALL subtypes, offering reliable diagnostic insights that can significantly support clinical decision-making. With the gene expression data of B-ALL patients as input, RanBALL is versatile and accepts various types of gene expression data as input, including raw counts, Fragments Per Kilobase of transcript per Million mapped reads (FPKM), and Transcripts Per Million (TPM). Different data types would be transformed into log_2_(TPM +1) for predicting the B-ALL subtypes. The processing pipeline encompassed four main steps: (1) data preprocessing and normalization, (2) RP-based dimension reduction, and (3) ensemble learning for multi-class classification, as depicted in Fig. 1C. This framework is particularly suited for the high-dimensional nature of gene expression data and the complex task of B-ALL subtyping, offering both accurate classification and improved visualization capabilities.

### Data preprocessing

The data preprocessing pipeline was illustrated as Fig. 1D. For the raw gene expression counts of 1988 B-ALL samples, only the gene expressed in at least 75% of the samples were retained, resulting in and finally 21,635 of the 52,007 original genes were kept. This filtering step eliminated rarely expressed genes that could introduce noise into the analysis. Subsequently, the raw read counts were then normalized to Transcripts Per Million (TPM), which allows for biological meaningful comparisons between different samples by adjusting for variations in the sequencing depth between samples (32). To prevent the inherent skewness of gene expression data, TPM values were transformed using the formula log_2_(TPM+1). Ultimately, the log-transformed TPM for subsequent model training and B-ALL subtype identification.

### Random projection

RP is a dimensionality reduction technique that aims to reduce the dimensionality of high-dimensional data while approximately preserving pairwise distances between data points. It is based on the Johnson–Lindenstrauss lemma (33) which provides a theoretical justification that a high-dimensional dataset can be approximately projected into a low-dimensional space while approximately preserving pairwise distances between data points. Specifically, the original *D*-dimensional data are projected onto a *d*-dimensional subspace through multiplying the original *D*-dimensional data matrix by the *d* x *N* random projection matrix. Namely,

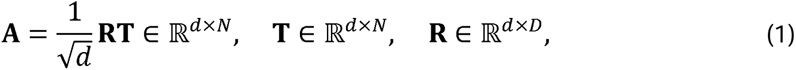

where **R** is random projection matrix, **T** is the original transcriptomic dataset, with *D* corresponding to the number of gene features and *N* denoting the number of B-ALL samples. The random projection matrix **R** should conform to any distributions with zero mean and unit variance, so that the random projection matrix **R** will give a mapping that satisfies the Johnson–Lindenstrauss lemma. For computational efficiency and the requirement of sparseness, we implemented a highly sparse RP method (34) This method determines the elements of **R** (i.e., *r_i,j_*) as follows:

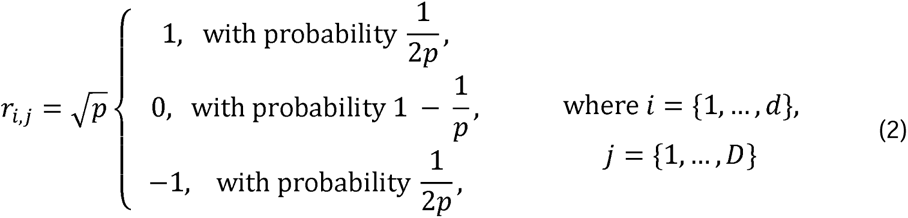

In accordance with the recommendation (34), we selected 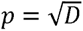.

### Ensemble learning model

To obtain reliable and robust performance, multiple RPs were applied into the original data matrix. The transformed low dimensional data matrix obtained from Eq. 1 was used for training an ensemble of SVM classifiers. To develop a robust model, we ensembled the predicted probability scores of each B-ALL subtype for low-dimensional data matrix and obtained an ensemble model. The ensemble score 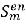 for each subtype was calculated by averaging all the prediction probability scores from each m-th SVM model in the ensemble:

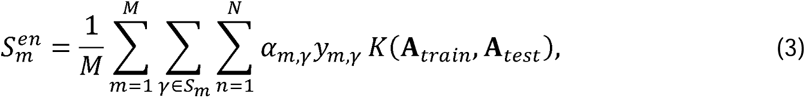

where S*_m_* is the set of support vector indexes corresponding to the *m*-th SVM, *M* is the ensemble size,*α_m,γ_* are the Lagrange multipliers, *N* is the number of predicted subtypes, *γ_m,γ_* is the class label for each subtype *K*(.,.) is the kernel function. Here, we used linear kernel for SVM after evaluating different kernel options. The **A***_train_* represents the projected RNA-seq data for training B-ALL samples, and the **A***_test_* corresponds to the test samples. To achieve optimal performance, we tried to optimize multiple hyperparameters including ensemble size (i.e., number of RPs to be used for ensemble), reduced dimensions, etc. Details can be found in the Results section below.

### RanBALL visualization

To capture both the global structure of the data and the specificity of subtype predictions, we developed a new visualization method for RanBALL (for ease of convenience, we refer to this as RanBALL visualization). Specifically, we utilized a weighted combination of two key matrices: a dimension-reduced feature matrix and a sample-to-subtype matrix derived from prediction results. The dimension-reduced feature matrix was obtained through RP techniques. This matrix was then normalized using Z-Score, centering and scaling the data along each dimension across samples. The prediction subtype for each sample was encoded by one-hot encoding to create a sample-to-subtype matrix, where each row corresponds to a sample, and each column represents a subtype. This matrix was then normalized using a Z-Score transformation across all samples to ensure that the data is centered and scaled, making the features comparable with the dimension-reduced matrix. These two matrices were then combined with different weights to formulate the final visualization matrix, combining the predicted subtype information with the dimensional features. We defined *w* as the weight ratio of the dimension-reduced feature matrix over the sample-to-subtype matrix. This weight can be adjusted to emphasize either the reduced feature space (*w* > 1) or the predicted subtype information (0 < *w* < 1) in the final visualization. This combined matrix served as the input for t-SNE visualization (35), allowing for a more informative and potentially more biologically relevant representation of the data. The predicted subtypes can provide additional information with the original gene expression profiling data for clustering data points in visualizations. Furthermore, the adjustable weight parameter provides flexibility for researchers to emphasize either global patterns or subtype-specific characteristics. By combining the global patterns from dimension-reduced feature matrix with the subtype-specific information from RanBALL predictions, the visualization provides a more faithful and informative representation of the data.

### Subtype-specific differential gene expression analysis

For subtype-specific differentially gene expression (DGE) analysis, the expected raw counts from B-ALL samples were used. Genes with low read counts were filtered out using a CPM cutoff threshold equivalent to a count of 10 reads. Normalization factors were then calculated using the TMM method (36), and the counts was further processed using the *voom* transformation. The normalized counts were analyzed with the *lmFit* and *eBayes* functions from the *limma* R package v3.54.2 (37). The cutoffs of FDR < 0.05, and |log_2_FC| > 1 were applied to define significantly differentially expressed genes. Heatmap plots were generated by Pheatmap package (1.0.12) (38).

### Performance evaluation

To evaluate model performance, we measured accuracy (*Acc*), F1-score (*F1*), and Matthews correlation coefficient (*MCC*) (39) as follows:

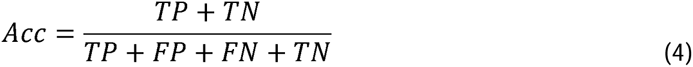

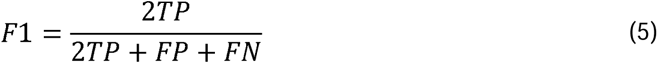

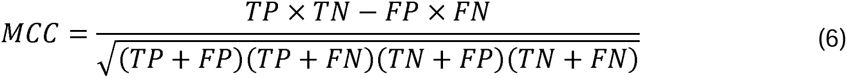

True positives (*TP*) denote the count of samples predicted to possess the specific subtype, which aligns with clinical documentation. False positives (*FP*) represent the count of samples incorrectly classified into different categories. True negatives (*TN*) indicate the count of samples predicted as ‘other’ that genuinely do not belong to the specified subtype category, while false negatives (*FN*) refer to the count of samples predicted as ‘other’ but are indeed found within the specified subtype category. The F1-score is a statistical measure used to evaluate the accuracy of a classification model, which is a way to balance the trade-off between precision and recall. A high precision might indicate a low tolerance for false positives, while a high recall might indicate a low tolerance for false negatives. The F1-score helps to find a balance between these two factors, making it a useful metric for evaluating the overall quality of a classification model. It is particularly useful in situations where the class distribution is imbalanced. In addition, MCC is a balanced measure that takes into account true and false positives and negatives. This makes it particularly helpful in imbalanced datasets where the number of positive instances may be very different from the number of negative instances.

## Results

### RanBALL preserves sample-to-sample distance

To explain the contributions of RP for dimension reduction in RanBALL, we investigated the degree of distortion caused by dimension reduction and compared the correlation of sample-to-sample distances after shrink with PCA (40), t-SNE (35) and UMAP (41), respectively, in different levels. We conducted Pearson correlation analysis to assess the similarities in sample-to-sample distances between the original and dimension-reduced data. As depicted in Fig. 2, RP achieved very high similarities in sample-to-sample distance, with the Pearson correlation coefficients exceeding 0.93. For example, when reducing the data to 1200 dimensions (from 21,635 to 1200), the correlation remained high at 0.94, indicating the preservation of almost all embedded information after dimension reduction. The remarkable performance of RP can be attributed to several key factors. One critical factor is RP’s ability to preserve pairwise distances (42), which plays a central role in maintaining high correlation coefficients between the original and projected data. This property is theoretically supported by the Johnson-Lindenstrauss lemma (34), which guarantees that a set of points in high-dimensional space can be projected onto a lower-dimensional space while approximately maintaining relative distances with high probability. Furthermore, RP’s linear transformation ensures that the overall structure of the data (43), including relative distances between samples, is preserved without introducing complex non-linear distortions. This simplicity not only enhances computational efficiency but also minimizes the risk of overfitting to specific data patterns. In contrast, correlations observed with PCA, t-SNE, and UMAP were notably lower (overall below 0.67, with a minimum of 0.32). This disparity in performance can be explained by the inherent characteristics of these methods. While effective for linear dimensionality reduction, PCA focuses on preserving directions of maximum variance, potentially losing information crucial for maintaining sample-to-sample distances but not significantly contributing to overall variance. As non-linear techniques designed for dimension reduction and low-dimensional visualization, t-SNE and UMAP focus on preserving local structure and often distort global structure. These could be the reasons to explain their poor performance in preserving overall sample-to-sample distances in this context. RP’s exceptional performance in preserving sample-to-sample distances while significantly reducing dimensionality makes it particularly well-suited for the high-dimensional, complex nature of gene expression data in B-ALL subtyping.

**Figure 2.**
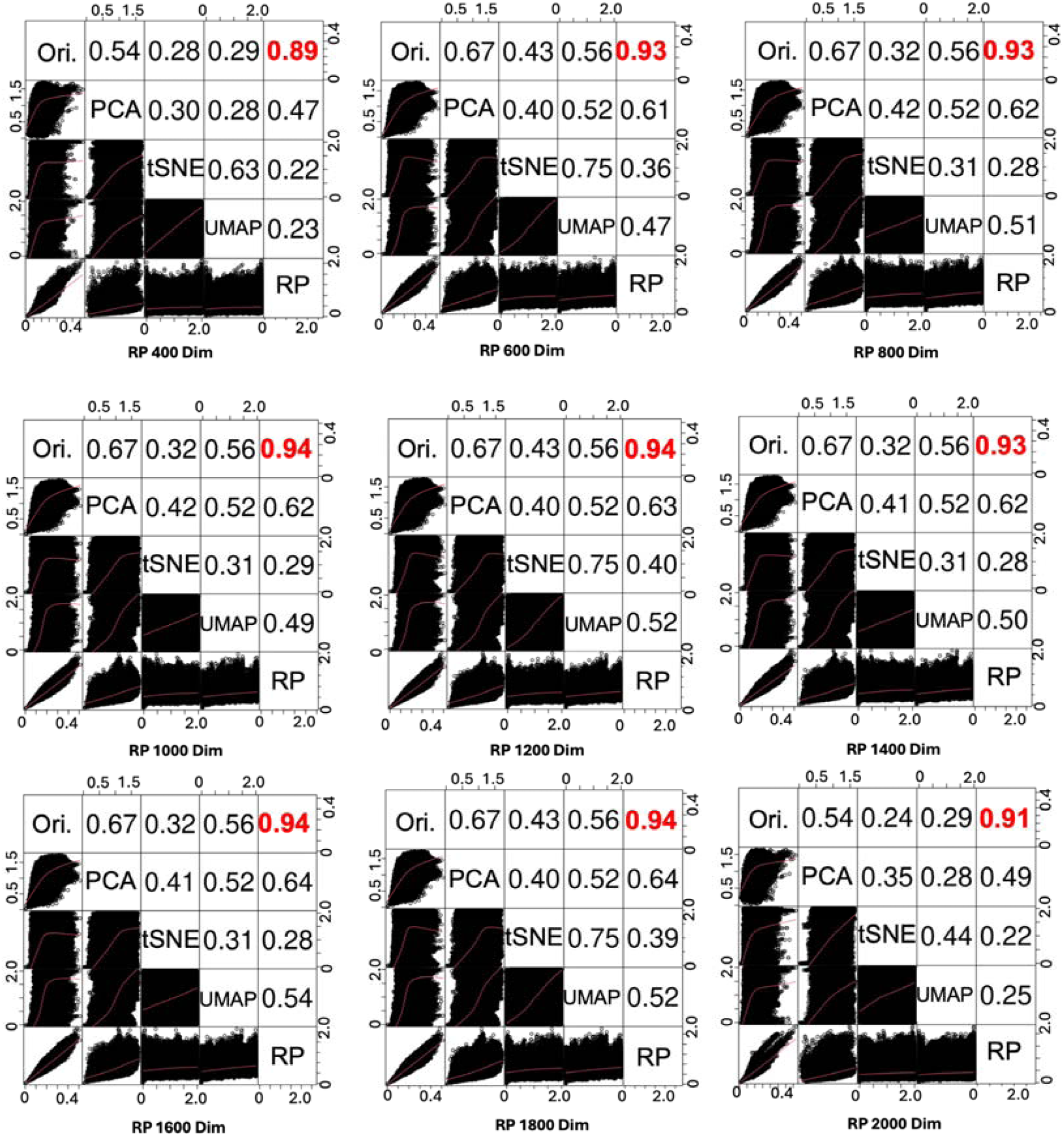
RP preserves sample-to-sample information better than state-of-the-art dimension reduction methods including PCA, t-SNE, and UMAP for RanBALL subtype identification. We compared RP with other state-of-the-art dimensionality reduction methods across different dimensions (400, 600,…, 2000). The upper triangular section of each matrix displayed the PCC between the sample-to-sample distances in the original high-dimensional space (Ori.) and the corresponding reduced-dimensional space for each method. Higher PCC values indicated better preservation of the original data structure. RP consistently achieved higher PCC (highlighted in red) that outperformed PCA, t-SNE, and UMAP. The lower triangular section provided scatter plots of pairwise distances between samples before and after dimensionality reduction, illustrating how well each method preserved the relative distances between points.

### The ensemble RP model performs better than individual RP models

To ensure the robust and stable performance of RanBALL, we applied ensemble learning to the predicted results obtained after dimensionality reduction with multi-class SVM. By aggregating predictions from multiple models, ensemble methods typically lead to better performance than relying on individual models. Additionally, ensemble methods help to reduce overfitting by averaging the biases of different models, thus providing a more generalizable solution. The original dimension (21,365) was reduced to a low dimension (we tried from 400 to 2000, with an interval of 200), to test the performances of different dimensions. Fig. 3A showed the performance of comparing ensemble models and individual models based on 100 runs of RPs. Focusing on overall accuracy metrics (Fig. 3A), the results revealed that the ensemble method’s prediction exhibited greater performance and stability with statistical significance compared to individual tests across all dimensions, indicating ensemble method’s superiority in generating stable and trustworthy prediction outcomes. To find the optimal reduced dimensionality, multiple dimensions of RP were applied in the model. The results showed that there was no significant difference for different dimensions when the number of dimensions is no less than 600 (Fig. 3B). The performance was stabilized when the dimensionality is no less than 1200. In addition, the performance variations of individual RP models with 100 times of running RP were limited when the dimensionality reached 1200 (Fig. 3B). Based on these results, 1200 was chosen for the subsequent model training. Next, we compared the performance with different ensemble sizes. Fig. 3C demonstrated that the ensemble size of 30 had more stable and robust performance in term of accuracy. Based on this finding, we selected the ensemble size of 30 for the model training. The empirical approach to determining these parameters ensures that the final model configuration is well-suited to the B-ALL subtyping with complex gene expression data.

**Figure 3.**
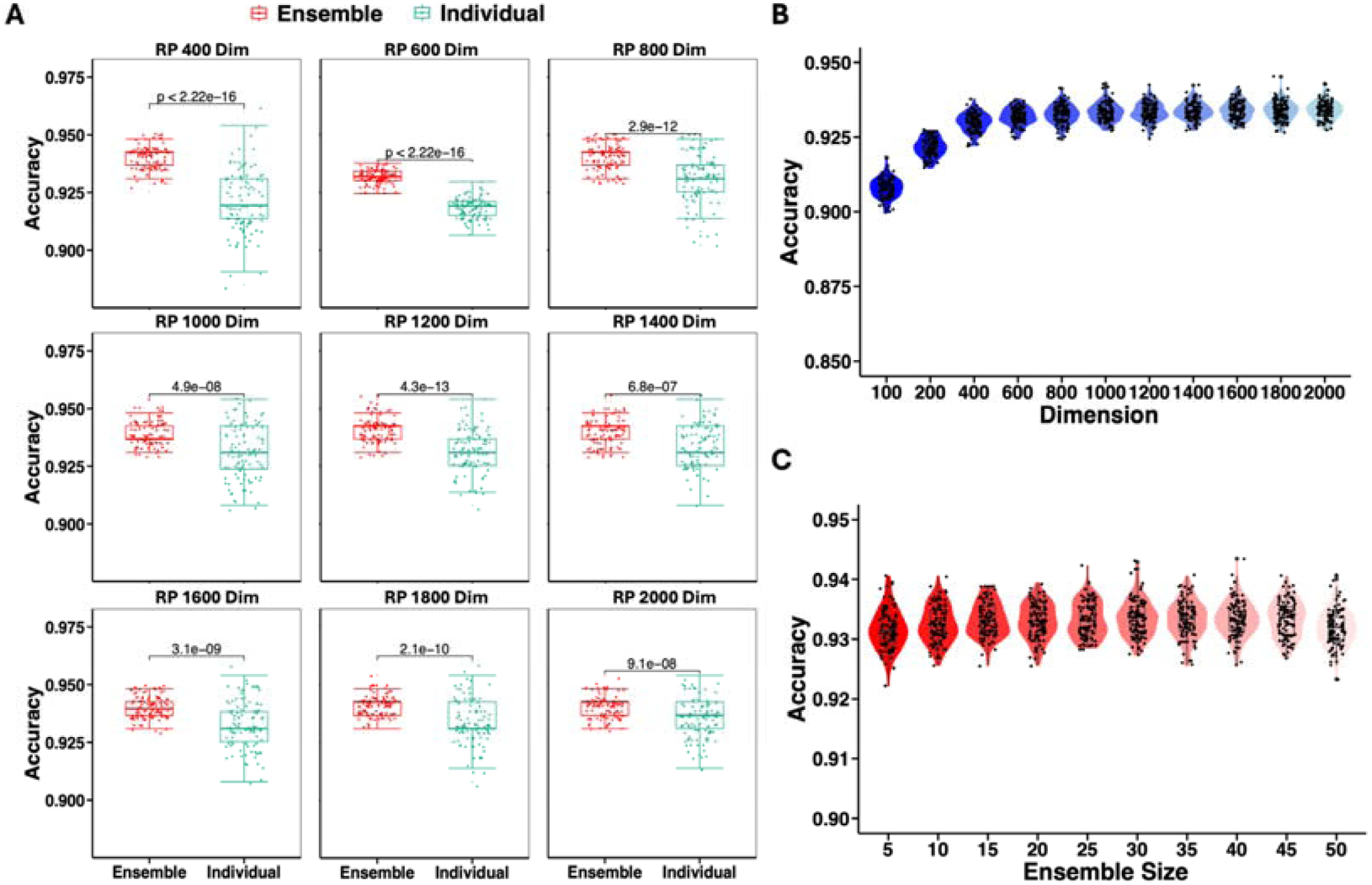
The performance of RanBALL in different RP models. **(A)** The ensemble RP model outperformed individual RP models across different reduced dimensions. Red boxes represented the accuracy distribution of the ensemble method aggregating 30 RPs, while green boxes denoted the accuracy distribution of individual classifiers on single RP. Statistical significance was assessed using the Wilcoxon signed-rank test, with p-values displayed above each comparison. **(B)** The model performance across different reduced dimensions. The violin plot illustrated the distribution of accuracy scores for dimensions ranging from 100 to 2000, with an interval of 200. **(C)** The model performance across different ensemble sizes. Violin plots depicted the distribution of accuracy scores for ensemble sizes ranging from 5 to 50. Black dots represented individual data points, while the violin shape showed the probability density of the data.

### RanBALL outperforms state-of-the-art methods for B-ALL subtyping

To assess the performance of the RanBALL model and its potential generalizability to unseen data, we employed a rigorous 100 times 5-fold cross-validation (CV) tests on an RNA-seq dataset comprising 1743 B-ALL samples with 20 subtypes as described in Fig. 1A. Our RanBALL model yielded impressive results exhibiting an accuracy of 93.35%, an F1-score of 93.10% and an MCC of 0.93 (Fig. 4A). These metrics collectively offered a comprehensive evaluation of the model’s efficacy. Given its exceptional performance across these metrics, the RanBALL model demonstrated significant promise for B-ALL subtyping. Additionally, we conducted a comparative analysis of the performance between RanBALL and ALLSorts (29), a well-established logistic regression classifier for B-ALL subtyping with the same data. As illustrated in Fig. 4A, RanBALL exhibited superior performance compared to ALLSorts in terms of Accuracy, F1-score and MCC. Notably, the superior F1-score of RanBALL suggested a more balanced trade-off between precision and recall relative to ALLSorts. The MCC performance matrix offered a balanced assessment even in scenarios where classes exhibit disparate sizes, indicating that RanBALL demonstrated superior performance particularly in multiclass classification settings with imbalanced class distributions compared to ALLSorts.

**Figure 4.**
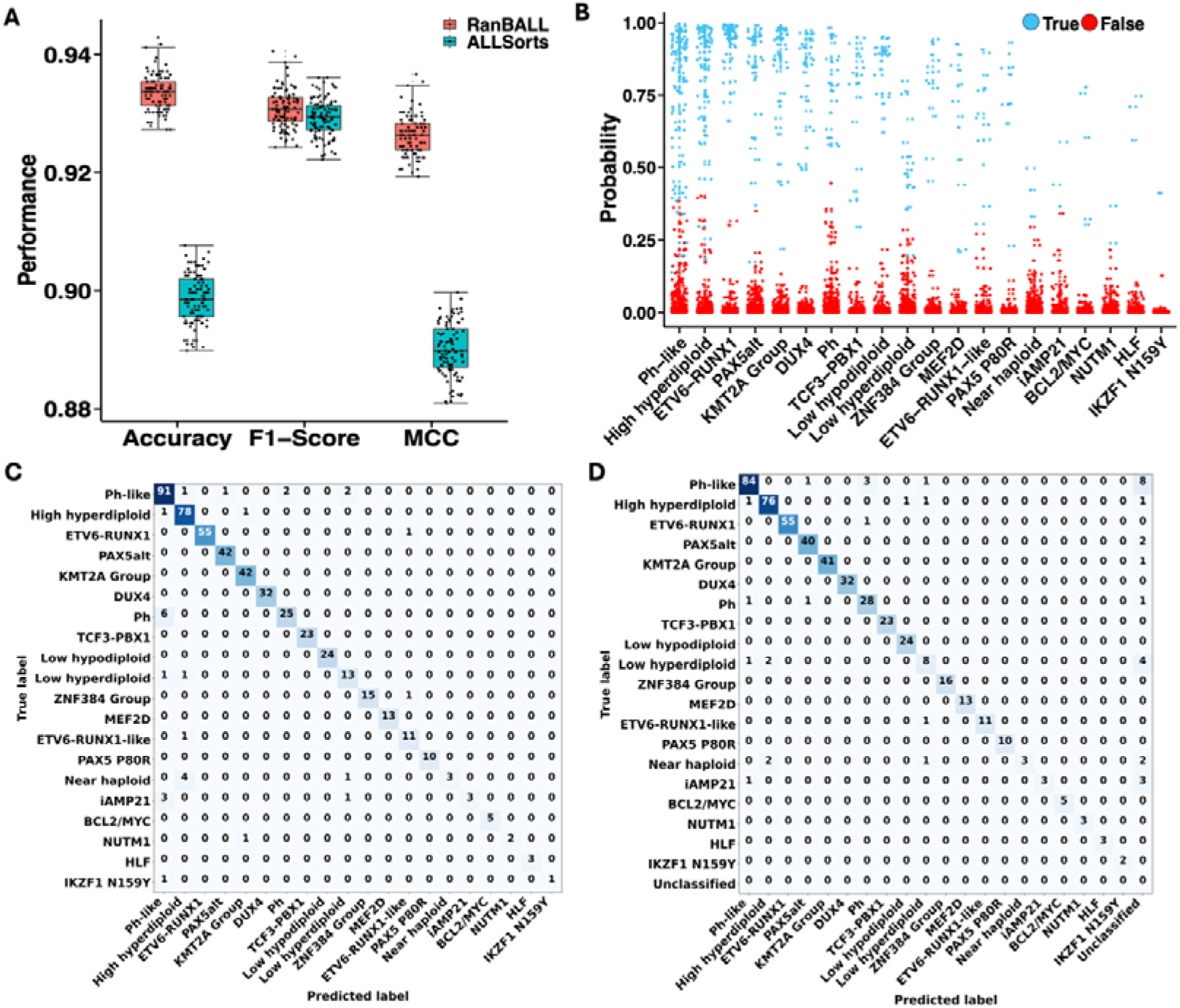
Comparing RanBALL with state-of-the-art methods like ALLSorts for B-ALL subtyping. **(A)** Comparing RanBALL and ALLSorts for identifying B-ALL subtypes in terms of various performance metrics. Accuracy, F1-score and MCC were used for evaluating model performance. Box plots illustrated the distribution of Accuracy, F1-score, and MCC across 100 times 5-fold cross validation. **(B)** Prediction probability distribution for the 30% held-out test set using RanBALL. Each point represents the probability of a sample (out of 521) being classified into a specific B-ALL subtype. Specifically, the blue dots indicate the specific subtype that the RanBALL model predicts to align with the categories on the horizontal axis. **(C, D)** Confusion matrices for the 30% held-out test set, comparing RanBALL (**C**) and ALLSorts (**D**) performance. Each element of the matrices shows the number of samples classified, with the diagonal representing correct classifications (True Positives). Color intensity correlates with the number of samples.

Subsequently, we applied the RanBALL model to hold-out tests to further demonstrate the superiority of RanBALL over state-of-the-art methods. Specifically, we selected one of the hold-out test sets, which was comprised of 521 samples, generated by randomly sampling 30% of the entire B-ALL dataset. The RanBALL model demonstrated a remarkable accuracy of 94.24% on this held-out test subset. The prediction probabilities of each test sample were shown in Fig. 4B, demonstrating the model’s consistent ability to maintain high confidence levels for accurate predictions. The robust performance of the model, evidenced by high-probability predictions, indicated its proficiency in distinguishing intrinsic data patterns, thereby yielding confident and reliable outcomes. Notably, it exhibited the capability to deliver accurate predictions even for subtypes characterized by limited sample sizes. However, it’s important to acknowledge that prediction probabilities for such subtypes may not attain exceptionally high levels. The 30% held-out test was also performed for ALLSorts, which achieved an accuracy of 89.64% on the same test dataset. The confusion matrices were illustrated in Fig. 4C-D, providing a detailed breakdown of the model’s prediction ability for each subtype in test data. Most significantly, RanBALL achieved superior classification accuracy for the Ph-like subtype, correctly identifying 91 out of 97 patients, surpassing ALLSorts’ performance of 84 correct classifications. Additionally, RanBALL showed improved performance in distinguishing Low hyperdiploid, accurately classifying 13 out of 15 cases, compared to ALLSorts’ 8 correct classifications. For some subtypes with similar characteristics and features, it is likely for the model to incorrectly predict the sample of one subtype to be another group. For example, 2 samples with Ph subtype were wrongly predicted in the Ph-like by RanBALL, while 3 were wrongly classified as the Ph subtype in the Ph-like by ALLSorts. This situation also occurred in the subtypes related to chromosome rearrangement (Near haploid, Low hyperdiploid, and High hyperdiploid), suggesting that B-ALL patients in these subtypes with similar characteristics and features were difficult to distinguish.

### RanBALL provides better data visualization compared to tSNE

RanBALL demonstrated superior visualization capabilities compared to traditional methods by incorporating predicted subtype information. We selected the t-SNE, one of the powerful and representative methods for visualizing high-dimensional data, to compare the performance of visualization. Fig. 5A illustrated the effectiveness of RanBALL in visualizing B-ALL samples, where the majority of subtypes were well-clustered, reflecting the model’s capability to maintain and highlight the inherent structure in the data. This visualization allowed for easy identification and interpretation of the 20 different subtypes, ranging from common subtypes like BCL2/MYC and DUX4 to rarer subtypes such as ZNF384 Group and iAMP21. Each subtype, represented by different colors and labels, formed tight, distinct clusters. In contrast, Fig. 5B presented a conventional t-SNE visualization without the integration of predicted subtype information, where subtype boundaries were less distinct and overlap more significantly. Subtypes such as High hyperdiploid, KMT2A Group, PH and Ph-like did not cluster as clearly, indicating that conventional tSNE was unable to distinguish subtypes of B-ALL patients with similar gene expression patterns. The difference between the visualization results by RanBALL and tSNE underscored the value of RanBALL’s approach in enhancing the interpretability and informative visual representations of complex transcriptomic data. By leveraging predicted subtype information, RanBALL not only improved visual clarity but also potentially revealed biologically meaningful relationships between subtypes. This enhanced visualization technique could provide valuable information for researchers in identifying patterns, outliers, and potential new subgroups within B-ALL samples, ultimately leading to better understanding and classification of this complex disease.

**Figure 5.**
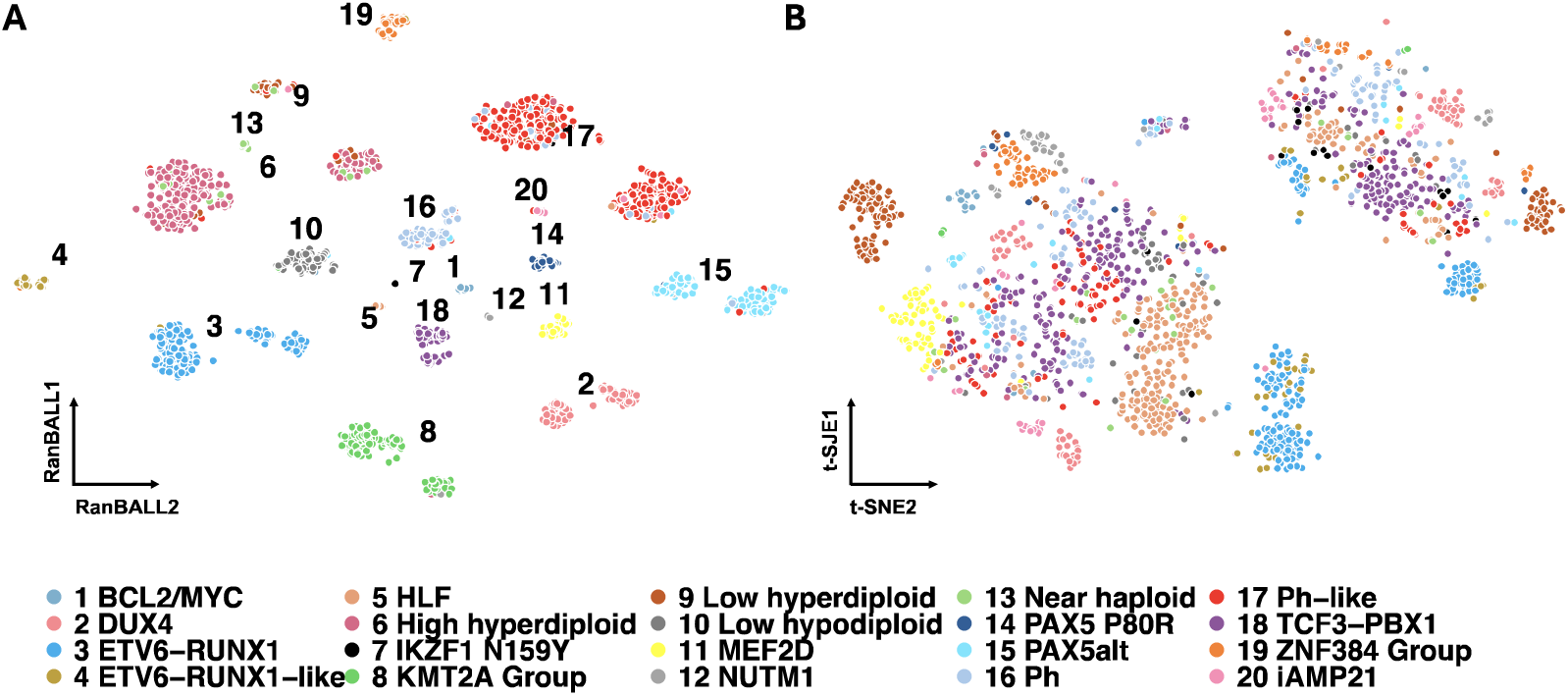
Comparing B-ALL and state-of-the-art visualization methods like tSNE for visualizing B-ALL subtype groups. **(A)** RanBALL visualization of the reduced dimension matrix incorporating predicted subtype information. **(B)** t-SNE visualization of the reduced dimension matrix with conventional gene expression profiling information only. The same color scheme was used in the two plots.

### Subtype-specific differential gene expression analysis of B-ALL patients

To identify and investigate the subtype-specific biomarkers for each B-ALL subtype, we performed subtype-specific differential gene expression analysis. Fig. 6A illustrated the differential expressed genes (DEGs) between Ph-like B-ALL and the rest subtypes. The expression plots of the upregulated DEG ENAM across all B-ALL samples were shown in Fig. 6C, highlighting its specific overexpression in the Ph-like subtype. The ENAM gene was specifically expressed at the samples with Ph-like subtype. The heatmap displayed the expression profiles of top 20 DEGs (Fig. 6B). It indicated the potential differences among subtypes within the biological functions and processes. Among the most upregulated genes, CRLF2, one of the most important genes in Ph-like ALL, was consistent with its known role in activating JAK-STAT signaling in a subset of Ph-like cases (44,45). Other significantly overexpressed genes, including GPR110, ENAM, LDB3, and IGJ, suggesting alterations in cell adhesion, signaling, and immunoglobulin production (44,46,47). Notably, SPATS2L overexpression has been associated with poor prognosis (48,49). We also conducted differential gene expression analysis on the PAX5alt subtype (Fig. 6D-F). These upregulated genes may play crucial roles in promoting cell proliferation, survival, and signaling pathways in PAX5alt B-ALL. For instance, TPBG was upregulated in high-risk cytogenetic subgroups and overexpressed on the plasma membrane of lymphoblasts collected at relapse in patients with B-cell precursor ALL (50). Similarly, KSR2, a kinase suppressor of Ras 2, has been implicated in dysregulation of multiple signaling (51), suggesting a similar altered signaling pathway in PAX5alt B-ALL. Additionally, TIFAB has been shown to regulate USP15-mediated p53 signaling in stressed and malignant hematopoiesis (52). Interestingly, NFATC4 significant upregulation in PAX5alt B-ALL contrasts with its significant downregulation in Ph-like B-ALL, highlighting distinct transcriptional programs between these subtypes. For differential gene expression analysis between High hyperdiploid and other subtypes (Fig. 6G∼I), the upregulated gene DDIT4L has been identified as therapeutic targets in PDX ALL carrying the recently described DUX4-IGH translocation (53). Notably, the upregulated gene OVCH2 was observed that it was downregulated in ALL (54,55). Additionally, S100A16 has been implicated in suppressing the growth and survival of leukemia cells in adults with Ph-negative B-ALL (56). The subtype-specific differential gene expression analysis for the remaining B-ALL subtypes were presented in Fig. S1-S6. These subtype-specific differential gene expression analyses revealed distinct molecular signatures and characteristics across B-ALL subtypes, which could expand our understanding of molecular mechanisms underlying different subtypes.

**Figure 6.**
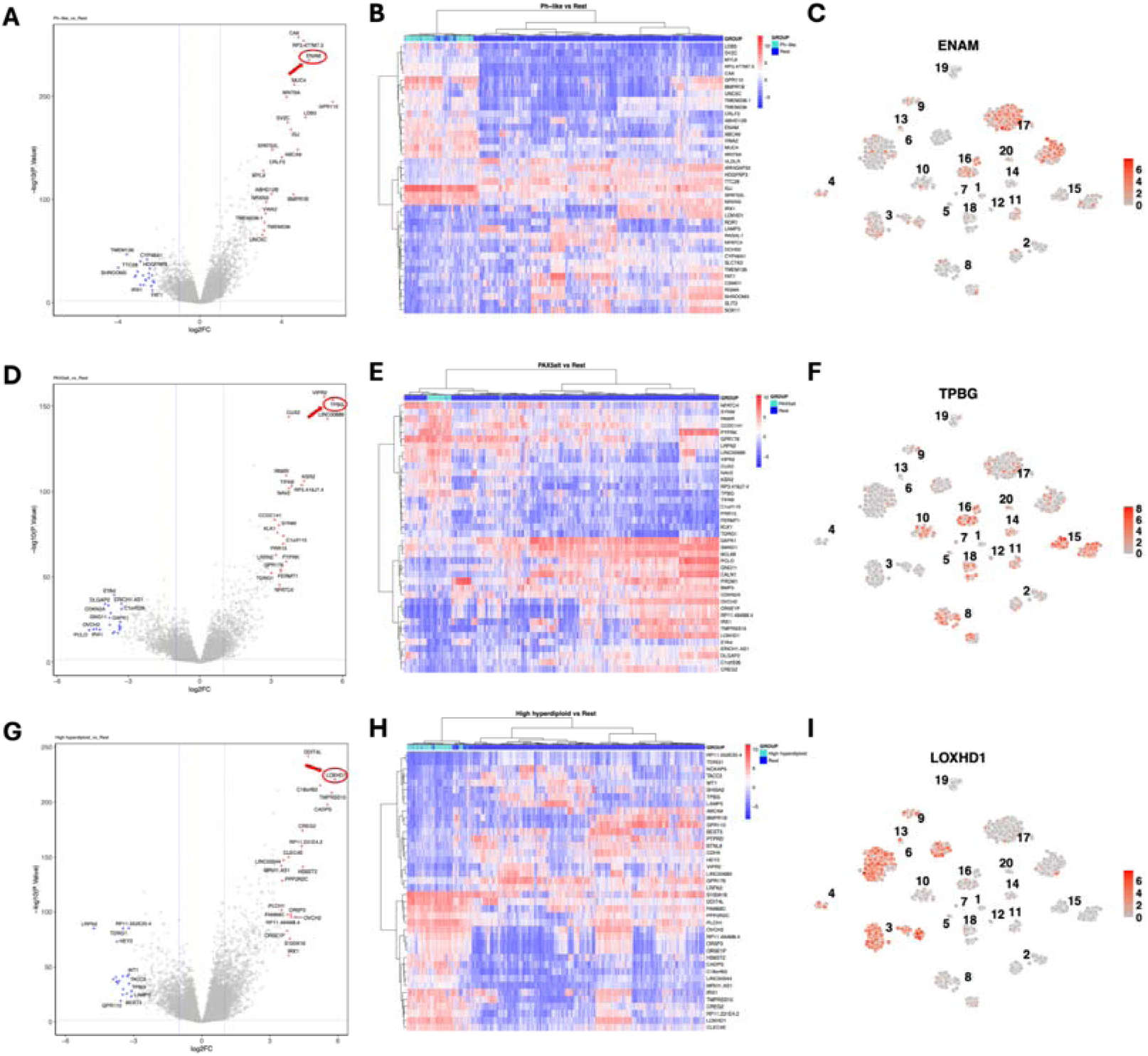
Subtype-specific differential gene expression analysis within B-ALL subtypes. (A, D, **G)** Volcano plots illustrated differential expression genes between specific B-ALL subtypes and all other subtypes. The x-axis represented log2 fold change, while the y-axis showed -log10(p-value). Red dots indicated 20 significantly up-regulated genes, blue dots represented 20 significantly down-regulated genes. Top 20 DEGs were labeled, with the most significant gene circled in red. **(A)** Ph-like vs. rest; **(D)** PAX5alt vs. rest; **(G)** High hyperdiploid vs. rest. **(B, E, H)** Heatmaps displayed expression patterns of the top 20 DEGs for each subtype comparison. Rows represented genes, columns represented samples. Color scale ranges from blue (low expression) to red (high expression). Hierarchical clustering dendrograms were shown for both genes and samples. Sidebar annotations indicated sample subtypes and relative level of gene expression. **(B)** Ph-like vs. rest; **(E)** PAX5alt vs. rest; **(H)** High hyperdiploid vs. rest. **(C, F, I)** The expression plot of the up-regulated DEG for Ph-like subtype. RanBALL plots visualized the expression levels of the significantly up-regulated gene for each subtype across all B-ALL samples. Each point represented a sample, colored by expression intensity (red: high, grey: low). Numbers indicated different B-ALL subtypes. **(C)** DEG for Ph-like (ENAM); **(F)** DEG for PAX5alt (TPBG); **(I)** DEG for High hyperdiploid (LOXHD1).

## Discussion

In this study, we introduced an ensemble-based model, RanBALL, which integrated ensemble RP and SVM techniques to accurately identify B-ALL subtypes using solely RNA-seq data. One of the key strengths of RanBALL lies in its ability to preserve sample-to-sample distances after dimensionality reduction, as evidenced by the high Pearson correlation coefficients (PCC) compared to PCA, t-SNE, and UMAP (Fig. 2). This preservation of data structure is crucial for accurate subtype classification and represents a significant advantage over traditional dimensionality reduction methods. Moreover, the integration of ensemble learning with RP enhanced the stability and reliability of predictions, mitigating the risks associated with imbalanced datasets and reducing overfitting. The experiments indicated that the ensemble learning method achieved superior stability and better performance than the individual RP models (Fig. 3A). In addition, RanBALL achieved remarkable performance metrics (accuracy: 93.35%, F1-score: 93.10%, MCC: 0.93) that significantly surpassed those of ALLSorts (Fig. 4A), one of the current state-of-the-art methods. This is particularly important given the heterogeneous nature of B-ALL, where accurate classification can directly influence risk stratification and treatment decisions. The high Moreover, MCC (92.62%) specifically indicated RanBALL’s effectiveness in handling the inherent class imbalance present in B-ALL subtypes, which is a common challenge in clinical datasets. The application of ML models in B-ALL subtype identification demonstrated the feasibility of leveraging complex datasets to discover subtle differences among patients. This approach overcame the limitations of traditional subtyping methods, which often rely on a limited set of markers and may not capture the full spectrum of disease heterogeneity.

Beyond classification, our enhanced visualization approach, which incorporates predicted subtype information, demonstrated better visualization capabilities compared to traditional methods. The incorporation of predicted subtype information into visualizations resulted in more distinct clusters, facilitating the discovery of novel subtype-specific markers. The subtype-specific differential gene expression analysis revealed distinct gene signatures that could serve as therapeutic targets or prognostic indicators across B-ALL subtypes, providing valuable insights into the translational research.

However, there is still room for improvement in certain B-ALL subtypes, necessitating further enhancement of prediction capabilities. First, future studies should aim to validate our models in more diverse and independent cohorts to ensure their broad applicability. Second, the predictive performance of our models could be influenced by technical and biological confounders (57), such as batch effects and sample quality. Rigorous data preprocessing and quality control measures will be essential to mitigate these factors in future work. Some common batch-effect correction methods, like ComBat (58), can be applied to mitigate the batch effects to better address real-world challenges and facilitate clinical applications (59). Additionally, the imbalance among B-ALL subtypes within the dataset may also potentially impede model performance. To address this issue, data augmentation techniques (60) can be applied to augment the representation of minority subtypes.

Furthermore, the integration of additional multi-omics and multi-modal data types, such as genetic (61,62), epigenetic (63,64) and imaging data (65,66), may potentially enhance the accuracy and reliability of ML models in B-ALL subtype identification. The advent of single cell sequencing technologies has revolutionized our ability to dissect heterogeneity of B-ALL, enabling the characterization of cellular subpopulations and their functional states at an unprecedented resolution. The integration of multi-scale multi-omics and multi-modality can provide valuable insights into the molecular landscape of B-ALL subtypes and inform personalized therapeutic approaches.

We anticipate that the deployment of RanBALL will yield direct positive impacts on clinical diagnosis, personalized treatment strategies, and risk stratification within the realm of biomedical research and clinical settings. This is particularly essential as distinct B-ALL subtypes may respond differentially to various treatments and are associated with diverse outcomes and survival rates. Moreover, precise subtype identification can aid clinicians in selecting the most effective treatment strategies for each patient. To facilitate further extending and accessibility of RanBALL, we have developed an open-source Python package, available at https://github.com/wan-mlab/RanBALL.

## Acknowledgements

The authors would like to express our gratitude to St. Jude Cloud platform (https://www.stjude.cloud), which provided publicly accessible genomic data. Special thanks to all members of Dr. Wan’s lab for insightful discussions. The abstract of this work was published at AACR Annual Meeting 2024 (67).

## Authors’ contributions

L.L.: data preprocessing, machine learning model development, data analysis and interpretation, manuscript preparation, editing, and review. H.X.: data analysis and interpretation, manuscript preparation, editing, and review. X.W.: manuscript preparation, editing, and review. Z.T.: manuscript editing and review. J.D.K.: manuscript editing and review. J.W.: manuscript editing and review. S.W.: study concept and design, manuscript editing and review.

## Data availability

The RNA-seq data of B-ALL samples can be publicly accessed from St. Jude Cloud (https://pecan.stjude.cloud/static/hg19/pan-all/BALL-1988S-HTSeq.zip).

## Code availability

The RanBALL package can be accessed at https://github.com/wan-mlab/RanBALL.

