## Supplementary Figure S1-S6 for "RanBALL: An Ensemble Random Projection Model for Identifying Subtypes of B-Cell Acute Lymphoblastic Leukemia"

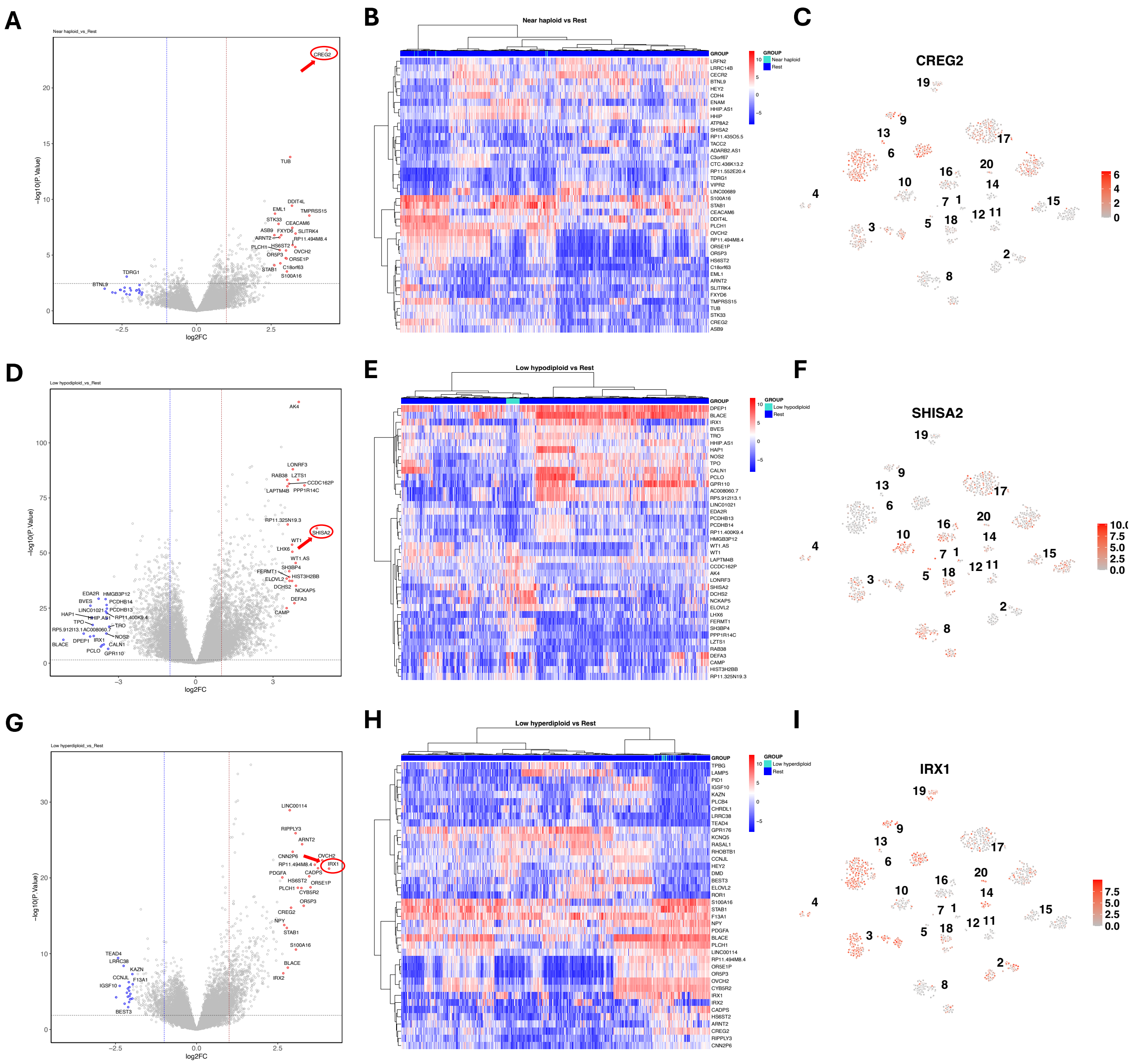


**Supplementary Figure S1. Subtype-specific differential gene expression analysis of Near haploid, Low hypodiploid, and Low hyperdiploid subtypes. (A, D, G)** Volcano plots illustrated differential expression genes between specific B-ALL subtypes and all other subtypes. **(A)** Near haploid vs. rest; **(D)** Low hypodiploid vs. rest; **(G)** Low hyperdiploid vs. rest. **(B, E, H)** Heatmaps displayed expression patterns of the top 20 DEGs for each subtype. **(B)** Near haploid vs. rest; **(E)** Low hypodiploid vs. rest; **(H)** Low hyperdiploid vs. rest. **(C, F, I)** The expression plot of the up-regulated DEG for specific B-ALL subtype. **(C)** DEG for Near haploid (CREG2); **(F)** DEG for Low hypodiploid (SHISA2); **(I)** DEG for Low hyperdiploid (IRX1).

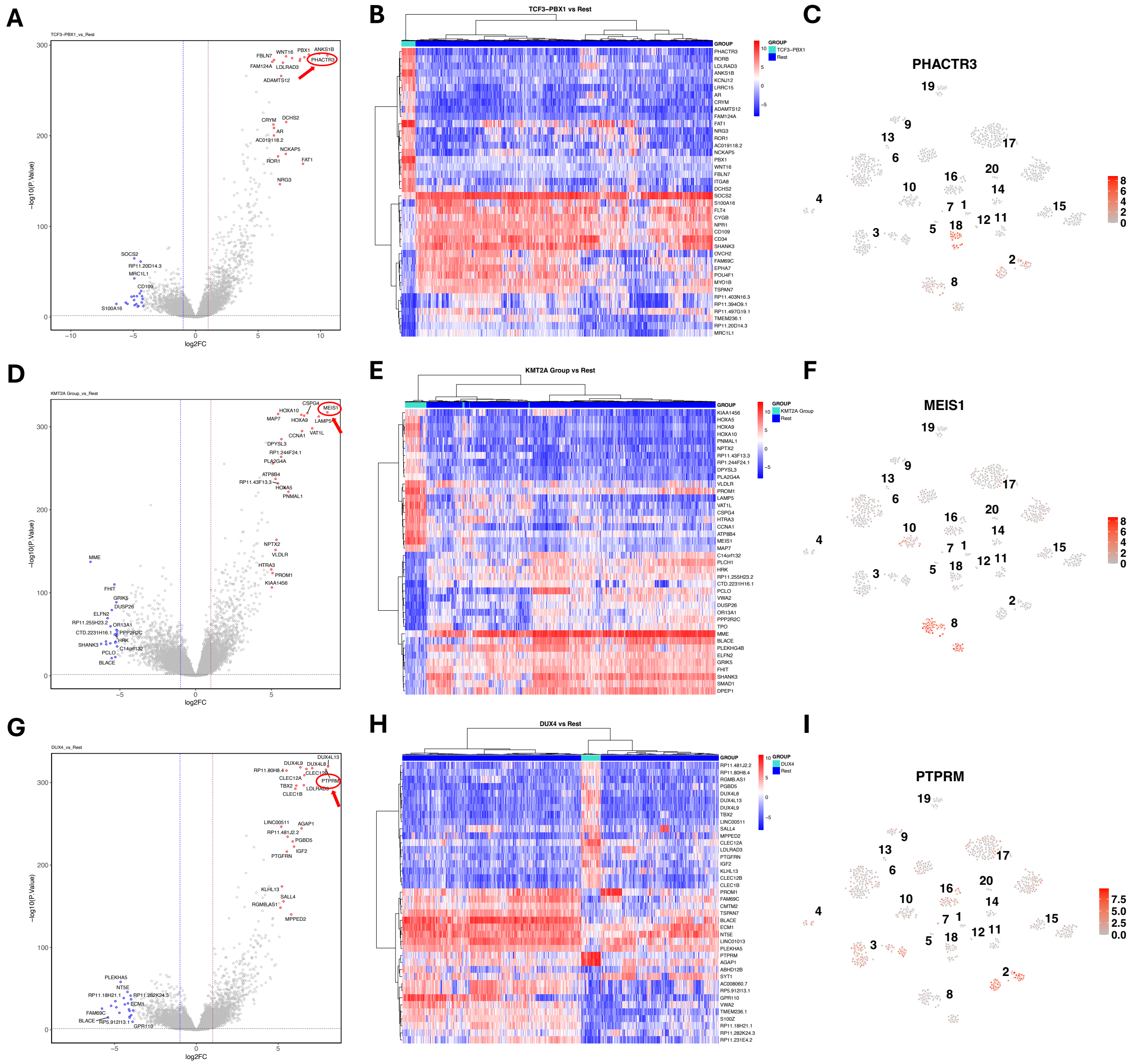

**Supplementary Figure S2. Subtype-specific differential gene expression analysis of TCF3-PBX1, KMT2A Group, and DUX4 subtypes. (A, D, G)** Volcano plots illustrated differential expression genes between specific B-ALL subtypes and all other subtypes. **(A)** TCF3-PBX1 vs. rest; **(D)** KMT2A Group vs. rest; **(G)** DUX4 vs. rest. **(B, E, H)** Heatmaps displayed expression patterns of the top 20 DEGs for each subtype. **(B)** TCF3-PBX1 vs. rest; **(E)** KMT2A Group vs. rest; **(H)** DUX4 vs. rest. **(C, F, I)** The expression plot of the up-regulated DEG for specific B-ALL subtype. **(C)** DEG for TCF3-PBX1 (PHACTR3); **(F)** DEG for KMT2A Group (MEIS1); **(I)** DEG for DUX4 (PTPRM).

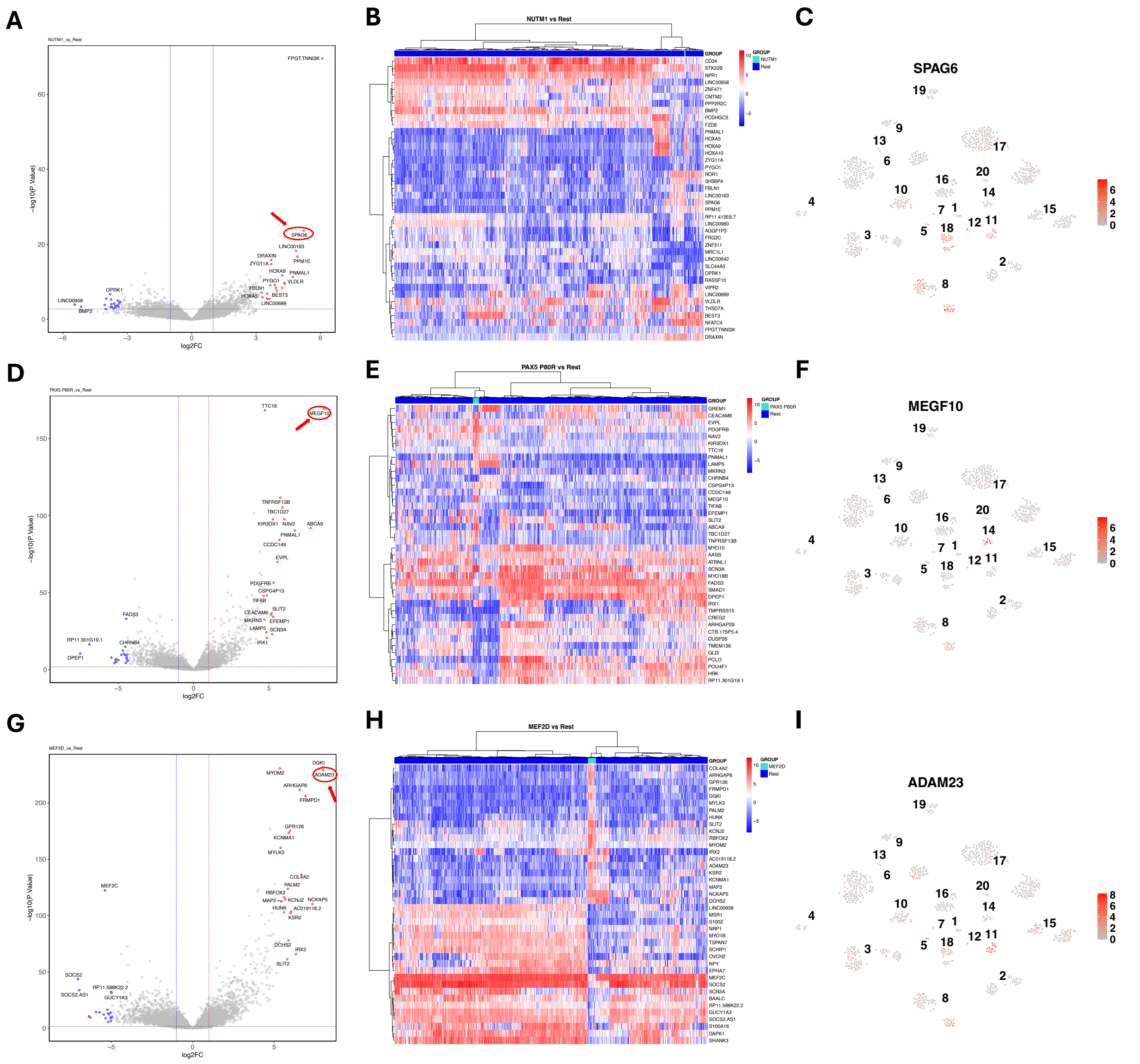

**Supplementary Figure S3. Subtype-specific differential gene expression analysis of NUTM1, PAX5 P80R, and MEF2D subtypes. (A, D, G)** Volcano plots illustrated differential expression genes between specific B-ALL subtypes and all other subtypes. **(A)** NUTM1 vs. rest; **(D)** PAX5 P80R vs. rest; **(G)** MEF2D vs. rest. **(B, E, H)** Heatmaps displayed expression patterns of the top 20 DEGs for each subtype. **(B)** NUTM1 vs. rest; **(E)** PAX5 P80R vs. rest; **(H)** MEF2D vs. rest. **(C, F, I)** The expression plot of the up-regulated DEG for specific B-ALL subtype. **(C)** DEG for NUTM1 (SPAG6); **(F)** DEG for PAX5 P80R (MEGF10); **(I)** DEG for MEF2D (ADAM23).

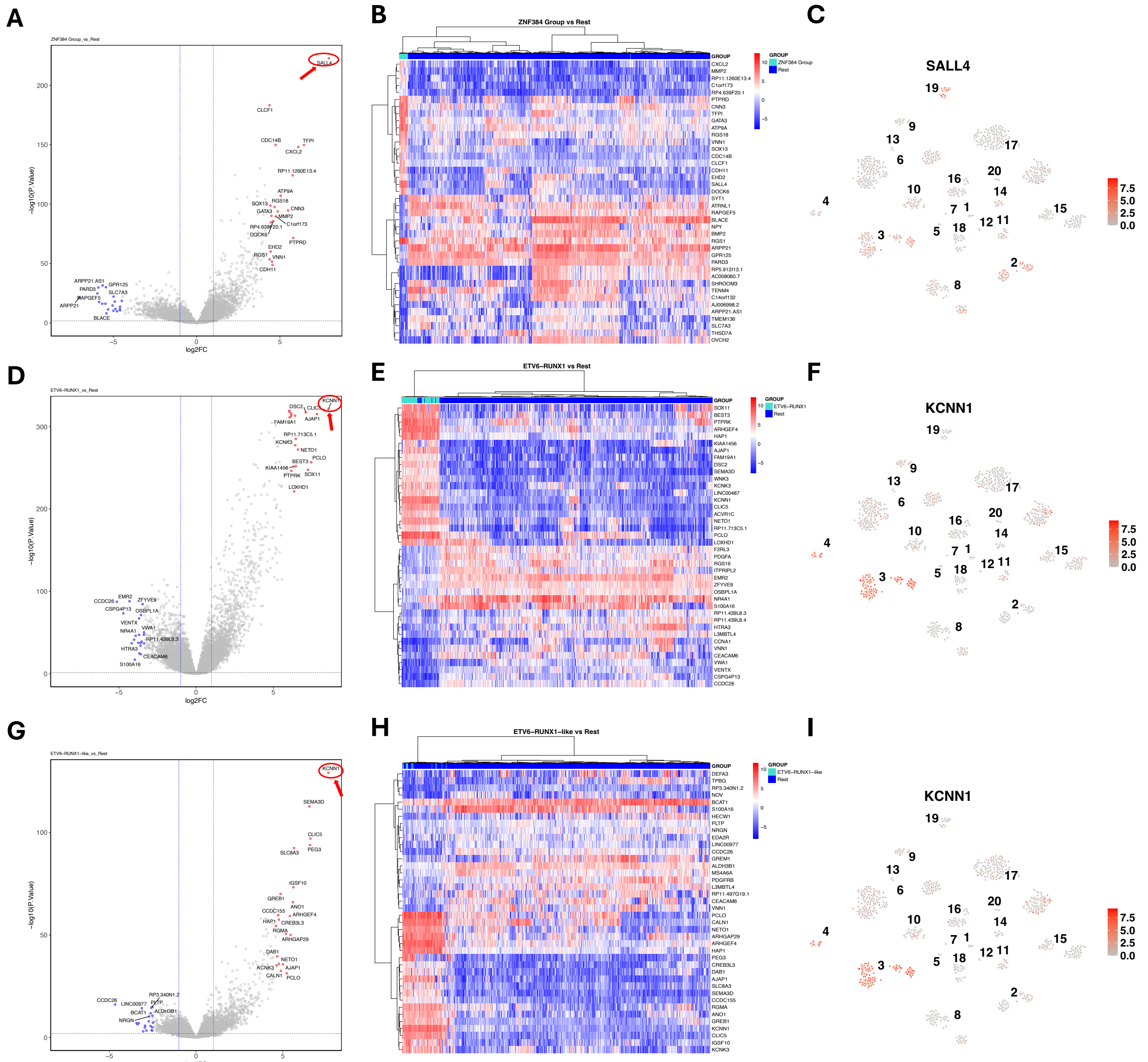

**Supplementary Figure S4. Subtype-specific differential gene expression analysis of ZNF384 Group, ETV6-RUNX1, and ETV6-RUNX1-like subtypes. (A, D, G)** Volcano plots illustrated differential expression genes between specific B-ALL subtypes and all other subtypes. **(A)** ZNF384 Group vs. rest; **(D)** ETV6-RUNX1 vs. rest; **(G)** ETV6-RUNX1-like vs. rest. **(B, E, H)** Heatmaps displayed expression patterns of the top 20 DEGs for each subtype. **(B)** ZNF384 Group vs. rest; **(E)** ETV6-RUNX1 vs. rest; **(H)** ETV6-RUNX1-like vs. rest. **(C, F, I)** The expression plot of the up-regulated DEG for specific B-ALL subtype. **(C)** DEG for ZNF384 Group (SALL4); **(F)** DEG for ETV6-RUNX1 (KCNN1); **(I)** DEG for ETV6-RUNX1-like (KCNN1).

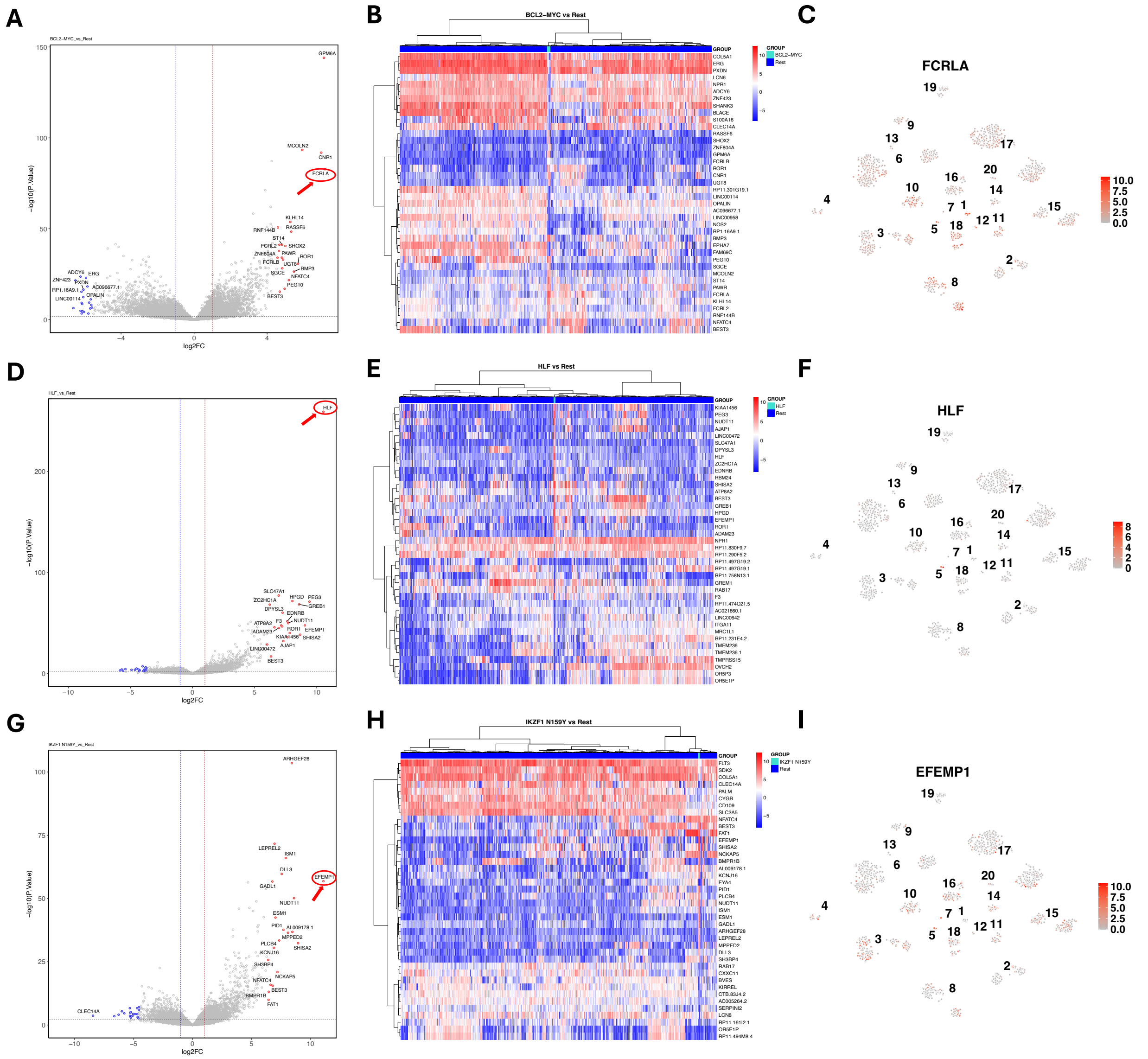


**Supplementary Figure S5. Subtype-specific differential gene expression analysis of BCL2/MYC, HLF, and IKZF1 N159Y subtypes. (A, D, G)** Volcano plots illustrated differential expression genes between specific B-ALL subtypes and all other subtypes. **(A)** BCL2/MYC vs. rest; **(D)** HLF vs. rest; **(G)** IKZF1 N159Y vs. rest. **(B, E, H)** Heatmaps displayed expression patterns of the top 20 DEGs for each subtype. **(B)** BCL2/MYC vs. rest; **(E)** HLF vs. rest; **(H)** IKZF1 N159Y vs. rest. **(C, F, I)** The expression plot of the up-regulated DEG for specific B-ALL subtype. **(C)** DEG for BCL2/MYC (FCRLA); **(F)** DEG for HLF (HLF); **(I)** DEG for IKZF1 N159Y (EFEMP1).

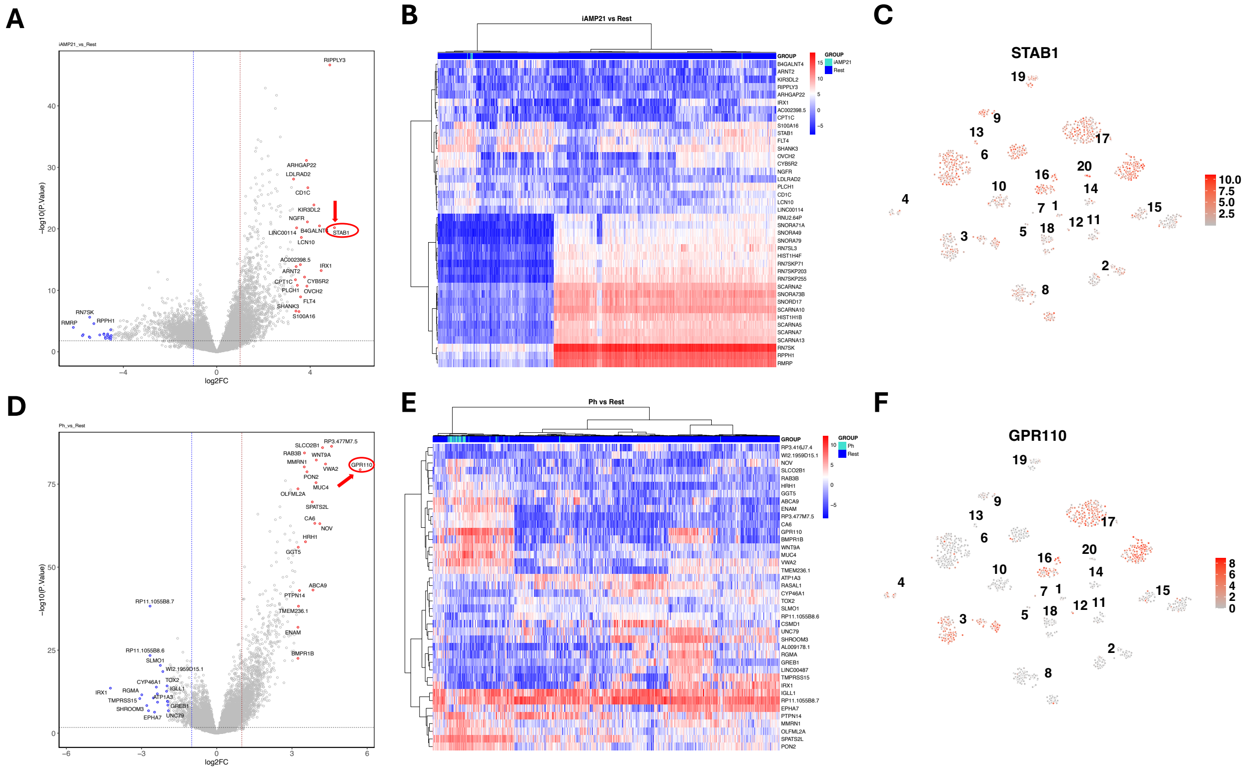

**Supplementary Figure S6. Subtype-specific differential gene expression analysis of iAMP21, and Ph subtypes. (A, D)** Volcano plots illustrated differential expression genes between specific B-ALL subtypes and all other subtypes. **(A)** iAMP21 vs. rest; **(D)** Ph vs. rest. **(B, E)** Heatmaps displayed expression patterns of the top 20 DEGs for each subtype. **(B)** iAMP21 vs. rest; **(E)** Ph vs. rest. **(C, F)** The expression plot of the up-regulated DEG for specific B-ALL subtype. **(C)** DEG for iAMP21 (STAB1); **(F)** DEG for Ph (GPR110).
